# The selective D3-Receptor antagonist VK4-116 effectively treats behavioral inflexibility in rats caused by self-administration and withdrawal from cocaine

**DOI:** 10.1101/2023.09.03.556083

**Authors:** Marios C. Panayi, Shohan Shetty, Micaela Porod, Lisette Bahena, Zheng-Xiong Xi, Amy Hauck Newman, Geoffrey Schoenbaum

## Abstract

Chronic psychostimulant use can cause long lasting changes to neural and cognitive function that persist even after long periods of abstinence. As cocaine users transition from drug use to abstinence, a parallel transition from hyperactivity to hypoactivity has been found in orbitofrontal-striatal glucose metabolism, and striatal D2/D3 receptor activity. Targeting these changes pharmacologically, using highly selective dopamine D3 receptor (D_3_R) antagonists and partial agonists, has shown significant promise in reducing drug-taking, and attenuating relapse in animal models of cocaine and opioid use disorder. However, much less attention has been focused on treating inflexible and potentially maladaptive non-drug behaviors following chronic psychostimulant use. Here we tested the selective D_3_R antagonist VK4-116 as a treatment for the long-term behavioral inflexibility in abstinent male and female rats with a prior history of chronic cocaine use. Rats were first trained to self-administer cocaine (0.75 mg/kg/reinforcer) or a sucrose liquid (10%, .04 mL/reinforcer) for 2 weeks (FR1 schedule, max 60 reinforcers in 3 hrs/ day), followed by 4 weeks of abstinence. Cognitive and behavioral flexibilities were then assessed using a sensory preconditioning (SPC) learning paradigm. Rats were given an VK4-116 (15 mg/kg, i.p.) or vehicle 30 mins prior to each SPC training session, thus creating four drug-treatment groups: sucrose-vehicle, sucrose-VK4-116, cocaine-vehicle, cocaine-VK4-116. The control groups (sucrose-vehicle, sucrose-VK4-116) demonstrated significant evidence of flexible SPC behavior, whereas cocaine use (cocaine-vehicle) disrupted SPC behavior. Remarkably, the D_3_R antagonist VK4-116 mitigated this cocaine deficit in the cocaine-VK4-116 group, demonstrating flexible SPC to levels comparable to the control groups. These preclinical findings demonstrate that highly selective dopamine D_3_R antagonists, particularly VK4-116, show significant promise as a pharmacological treatment for the long-term negative behavioral consequences of cocaine use disorder.

## Introduction

Substance use disorders (SUDs) are characterized by persistent drug use despite negative consequences and repeated intentions to abstain (Frazer et al., 2018; Spronk et al., 2013). In drug users, including cocaine use disorder (CUD), this behavioral inflexibility can reflect a lack of insight (Goldstein et al., 2009; Hester et al., 2007; Moeller et al., 2014). A lack of insight has been operationalized as an inability to mentally simulate and infer the causes and effects of one’s own behavior (Lucantonio et al., 2014; Raftery et al., 2020) and is associated with reduced function in orbitofrontal cortex (OFC) networks (Goldstein et al., 2009; Hart et al., 2020; Jones et al., 2012). Consistent with this, chronic cocaine use in CUD is consistently associated with decreased functional activity of the orbitofrontal cortex (OFC), and in a preclinical model, abstinent rats with a history of cocaine self-administration (SA) are impaired on insight-based behavioral tasks that require integrating learned information in the OFC to make inferences e.g. if A->B and B->C, then A->C (Lucantonio et al., 2015; Wied et al., 2013). These deficits are associated with inflexible task representations in OFC and can be successfully treated by optogenetic activation of pyramidal neurons in OFC (Lucantonio et al., 2014). This suggests that targeting OFC networks is a possible therapeutic pathway to treat the long-term consequences of chronic cocaine use.

Cocaine use has also been associated with increased D_3_R availability in the striatum and midbrain (Boileau et al., 2012; Matuskey et al., 2014; Payer et al., 2014; Segal et al., 1997; Staley et al., 1994; Staley & Mash, 1996), an increase which correlates with OFC hypofunction in CUD patients (Volkow et al., 1991, 1992, 1993), as well as in rodent and non-human primate models (Groman et al., 2016, 2020). Midbrain D_3_Rs in substantia nigra and ventral tegmental area are thought to be inhibitory modulators of presynaptic dopamine release in striatum and OFC (Gurevich & Joyce, 1999; Heidbreder & Newman, 2010; Koob & Volkow, 2010; Newman et al., 2021; Sesack & Grace, 2010). In non-clinical populations, midbrain D_3_R availability is correlated with both resting state OFC activity, and the functional connectivity between OFC subregions and multiple key large-scale neural systems such as salience executive control networks, basal ganglia/limbic network, and the default mode network (Cole et al., 2012). Recent evidence from cocaine self-administration and forced abstinence studies in rodents and non-human primates also supports a causal link between cocaine use, increased midbrain D_3_R availability, OFC hypofunction, and behavioral deficits in tasks like probabilistic reversal learning (Groman et al., 2016, 2020; Jedema et al., 2021). Indeed, D_3_R-antagonists/partial agonists have been shown to successfully reduce both self-administration and relapse to drug seeking as assessed by reinstatement to drug seeking caused by psychostimulants such as cocaine and nicotine in rodents (Jordan et al., 2020; Ross et al., 2007; Z.-X. Xi et al., 2005; Z. X. Xi et al., 2006), as well as opioids such as oxycodone and heroin (Boateng et al., 2015; Galaj et al., 2022; Jordan, Humburg, Rice, et al., 2019; Newman et al., 2023; You, Bi, Galaj, Kumar, Cao, Gadiano, Rais, Slusher, Gardner, Xi, & Newman, 2019). Therefore, it might be possible to treat the cognitive and behavioral sequalae of chronic cocaine use pharmacologically with D_3_R-antagonists that directly target increased D_3_R availability to indirectly target OFC hypofunction (Galaj et al., 2020; Groman et al., 2020; Newman et al., 2021).

Until recently, most of the available D_3_R-antagonists have had only moderate (<100-fold) D_3_R/D_2_R selectivity (Newman et al., 2021). However, this limitation has been overcome by the development of class of highly selective D_3_R-antagonists with low D_2_R binding affinity, including the compound VK4-116 (Jordan, Humburg, Thorndike, et al., 2019; Kumar et al., 2016; You, Bi, Galaj, Kumar, Cao, Gadiano, Rais, Slusher, Gardner, Xi, Newman, et al., 2019). Here we tested whether the novel D_3_R-antagonist VK4-116 could treat the deficits in insight-based behavior that occur following chronic cocaine use in a preclinical rodent model (Lucantonio et al., 2014, 2015; Wied et al., 2013). Rats were first trained to either self-administer cocaine (Coc) or sucrose (Suc), followed by 4 weeks of forced withdrawal. Next, rats were treated with either the D_3_R-antagonist VK4-116 (D3a) or vehicle (Veh) immediately prior to training each day on sensory preconditioning (SPC), a task that measures behavioral inference (Kahnt & Schoenbaum, 2021). SPC provides a measure of whether subjects make inferences about the relationships between cues and outcomes; first the cue-cue relationships A->B and C->D, and then the cue-outcome relationships B->Reward and D->Nothing. In a final test, responding more to cue A than C in anticipation of reward is thought to reflect the successful inference that A->B->Reward and C->D->Nothing (Killcross & Blundell, 2002; Rizley & Rescorla, 1972). Consistent with earlier reports, cocaine (Coc_Veh), but not sucrose (Suc_Veh), use disrupted evidence of behavioral inference in SPC in vehicle treated rats (Wied et al., 2013). However, both cocaine (Coc_D3a) and sucrose (Suc_D3a) treated rats showed intact behavioral inference in SPC, suggesting that the D_3_R-antagonist effectively treated the impairment in behavioral inference caused by cocaine-experience. Importantly, these effects were restricted to behavioral inferences in the final test phase and did not reflect differences in behavior or learning from earlier task stages. A detailed analysis of individual differences in behavioral patterns revealed additional support for this the efficacy of the D_3_R-antagonist treatment. These findings provide the first preclinical evidence for the efficacy of the selective D_3_R-antagonist VK4-116 as a pharmacological treatment for impaired insight and behavioral inflexibility following cocaine use and CUD.

## Materials and Methods

### Subjects

Subjects were 45 female and 45 male (N = 90) Long-Evans rats (250-300g, ∼3 months old) (Charles-River Laboratories, Wilmington, MA, USA) housed individually upon arrival, maintained on a 12hr light:dark cycle, and tested during the light phase. Rats were given at least 7 days to acclimate to the animal facility with *ad-libitum* access to food and water prior to any experimental procedures. Rats were food restricted 3 days prior to behavioural testing and pre-exposed to the relevant food reinforcer (10% w/v sucrose liquid and Banana flavored sucrose pellets (Bio-Serv, Frenchtown, NJ), self-administration and sensory preconditioning training respectively), but returned to *ad-libitum*food access during the forced cocaine withdrawal period. During food restriction, rats were fed at the end of each testing day (approximately 10 or 15g per female and male rat, respectively), and weights were maintained between above 85% of their baseline body weight. The experiment was conducted in 5 separate cohorts run sequentially: Cohort 1 (n = 6 Female, n = 6 Male), Cohort 2 (n = 6 Female, n = 6 Male), Cohort 3 (n = 6 Female, 6 Male), Cohort 4 (n = 12 Female, n = 12 Male), Cohort 5 (n = 15 Female, 15 Male). Experiments were performed at the National Institute on Drug Abuse, Intramural Research Program (NIDA-IRP), in accordance with the US National Institutes of Health (NIH) guidelines and were approved by the NIDA-IRP Animal Care and Use Committee (ACUC).

### Apparatus

Self-administration was conducted in standard modular rodent behavior boxes (Cohorts 1-3: Med Associates Inc., Georgia, VT, USA; Cohorts 4-5: Coulbourn Instruments, Allentown, PA, USA) inside sound attenuating chambers. Self-administration boxes were equipped with a retractable lever and a non-retractable lever on the right and left side of the front wall (designated as the active and inactive lever, respectively), and a house light centered at the top of the back wall. For cocaine self-administration, the back mounted catheter harness was connected to silastic tubing encased in a metal spring that was mounted to a swivel arm above the operant box and connected to an infusion pump mounted in the sound attenuating chamber (Instech Laboratories, PA, USA). For sucrose self-administration the boxes were equipped with a recessed magazine located between the two levers that provided access to 0.04 mL of sucrose liquid (10% w/v) via a retractable dipper cup.

Sensory preconditioning training was conducted using the same type of standard rodent behavior boxes (Coulbourn Instruments, Allentown, PA, USA) in sound attenuating chambers but located in a separate room. These boxes were equipped with a recessed food magazine in the center of the front wall, connected to a pellet dispenser mounted on the outside of the box that could deliver 45 mg banana flavored sucrose pellets (F0024, Bio-Serv, NJ, USA) into a food cup at the base of the magazine. Auditory stimuli could be delivered via a clicker mounted to the top-right of the front wall, a speaker on the opposite (top-left) of the back wall connected to a white-noise generator, and a speaker mounted on the top-right of the back wall connected to a tone generator. The auditory stimuli were a clicker (2 Hz), white noise (∼70 dB), a steady tone (1500 Hz, ∼75 dB), or a siren (oscillating at 5 Hz between 1000 Hz and 2000 Hz tones, ∼75 dB).

### Surgical procedures

Prior to behavioural testing, chronic jugular catheters were implanted under aseptic conditions (Mueller et al., 2021) . Rats were anaesthetized with either ketamine (100 mg/kg, i.p., Sigma) and xylazine (10 mg/kg, i.p., Sigma) (Cohorts 1-3), or isoflurane gas (induction at 4% and maintained at 1-2%, 2L/min O_2_) (Cohorts 4-5). A 22-gauge catheter (part# C30PU-RJV1412, Instech Laboratories, PA, USA) was inserted into the jugular vein and passed through an incision on the rat’s back to connect to a vascular access harness (part# VAH95AB14, Instech Laboratories, PA, USA). Some rats assigned to the sucrose self-administration condition received sham surgery in which the jugular vein was exposed but no catheter was implanted. Rats recovered for at least 7 days prior to self-administration training. Catheters were flushed daily with 0. 1 ml of mixture solution of gentamycin (1 mg/ml) and heparinized (1000 unit/ml)saline to maintain patency. Prior to self-administration training, catheter patency was assessed by observing loss of muscle tone following a brief methohexital (brevital) challenge (0.1 ml of 10 mg/ml brevital, i.v. via the catheter). Rats with patent catheters were then assigned to the cocaine self-administration groups, and the remaining rats with patent/non-patent catheters or sham surgeries were assigned to the sucrose self-administration groups.

### Drugs

Cocaine-HCL (NIDA IRP, Baltimore, MD, USA) was dissolved in saline, and administered through i.v. catheters at a concentration of 0.75 mg/kg/infusion (0.1 ml per infusion). To account for the range of rat weights during self-administration training, rats were weighed each day and assigned to a 20g weight interval (e.g. 300 −320g), and received a cocaine-HCL concentration corresponding to the average of the weight band (e.g. 310g). Weights remained relatively stable, and the rats did not need to change cocaine-HCL concentrations during self-administration training.

VK4-116 [(±)-N-(4-(4-(3-chloro-5-ethyl-2-methoxyphenyl)piperazin-1-yl)-3-hydroxybutyl)-1H-indole-2-carboxamide] was synthesized using the published method (Kumar et al., 2016), and dissolved in vehicle to be administered at a dose of 15 mg/kg. Vehicle was 25% (w/v) 2-hydroxypropyl-β-cyclodextrin (vendor#: 332607; Sigma-Aldrich St. Louis, MO, USA) dissolved in distilled water. VK4-116 and vehicle injections were administered i.p. at a volume of 1 ml/kg. All drugs were prepared fresh and kept for up to five days refrigerated.

### Self-administration (SA)

One day prior to SA, rats assigned to the sucrose condition were given one pretraining session with two phases to familiarize them with the sucrose dipper cups. During phase one (1 hr) the sucrose dipper was presented non-contingently for 40s approximately every 80s (variable inter-reinforcer interval schedule). During phase two (1 hr) the active lever was inserted into the chamber for a maximum of 30 trials in which an active lever press resulted in lever retraction, 40s sucrose dipper presentation, and an 80s inter-trial interval.

Following this, rats were trained to self-administer intravenous cocaine-HCl (0.75 mg/kg/infusion, 4s per infusion) or oral sucrose (0.05 ml, 10%, w/v) under a fixed ratio 1 schedule (FR1), in a 3-hr session/day for 14 consecutive days. Each session consisted of trials where the active lever was inserted, signaled the availability of a reinforcer. A single response on the active lever (FR1) resulted in the retraction of the active lever, and either a 4s presentation of the sucrose dipper (0.05 mL sucrose 10% w/v) or a 4s drug infusion (0.75 mg/kg). This was followed by a 40s timeout period (inter-trial interval) where the active lever was kept retracted and no reinforcers were available. A non-reinforced lever was always extended and present throughout the entire session, and responding on this lever was recorded but had no programmed consequences. Within each hour, rats could earn a maximum of 20 reinforcers and had at least 15 mins timeout with no reinforcers i.e. up to 45 min to complete 20 trials. This procedure ensured that cocaine rats received a maximum of 20 infusions per hour and 60 infusions per session/day to prevent overdose. Due to experimenter error, one cohort of rats (Cohort 1) in the sucrose SA condition could earn up to 100 reinforcers per hour.

### Withdrawal period

Following the completion of SA training, all rats were returned to *ad libitum* food access in their home cages with no experimental training for least 4 weeks of the withdrawal. Catheters and harnesses were removed at the start of the withdrawal period by cleaning and cutting the tubing such that it was no longer exposed. All rats received prophylactic treatment of enrofloxacin (0.454%, 1 ml, s.c.) for three days following catheter removal.

### Treatment

Rats in the sucrose (SA) and cocaine (Coc) SA conditions were pseudo-randomly allocated to either a vehicle (Veh) or D_3_R antagonist (D3a) treatment condition to create a total of four groups: Suc_Veh, Coc_Veh, Suc_D3a, and Coc_D3a. Pseudo-random allocation was used to roughly match the rates of SA behavior in each group, and equate the number of males and females. Treatment involved an i.p. injection of VK4-116 (15 mg/kg; D3a) or vehicle (25% β-cyclodextrin; Veh) on every day of the sensory preconditioning training. Rats were injected and returned to their home cage for 30 mins before being transferred to the behavioral training chamber for sensory preconditioning.

### Sensory preconditioning (SPC)

Sensory preconditioning (SPC) training began after the SA training and withdrawal period. The SPC procedure consisted of three stages, trained over 11 consecutive days, with drug treatment injections (Veh or D3a, 30 min pre-training) administered each day. Generally, rats received one session per day where each session contained 12 trials (variable inter-trial interval, M = 600s ± 300s), in which stimuli were presented non-contingently with or without reinforcers. The stimuli were four distinct 10s auditory cues, A and C (clicker and white noise, counterbalanced), B and D (tone and siren, counterbalanced). During conditioning, reinforcement entailed the delivery of 3 sucrose pellets within the 10s presentation of stimulus B, such that a pellet was delivered at 3, 6.5, and 10s. Immediately prior to preconditioning, rats were first shaped to receive food pellets from the magazine in a single session with 16 reinforcers (two sucrose pellets) delivered on a variable time schedule (*M* = 120s ± 60s).

1. Preconditioning: Rats received two days of preconditioning where two distinct S_1_-S_2_ stimulus pairs were presented: A->B and C->D. A stimulus pair consisted of a 10s auditory stimulus (clicker or white noise; A or C, counterbalanced) immediately followed by a second 10s auditory stimulus (tone or siren; B or D, counterbalanced). Each session consisted of a 6-trial block with one of the stimulus pairs (e.g. A->B), followed by a second block of the other stimulus pair (e.g. C->D). Block order was reversed on the second session, and fully counterbalanced across rats.
2. Conditioning: After 2 days of preconditioning, rats received 6 days of Pavlovian conditioning. Each session involved six reinforced trials of cue B (3 sucrose pellets delivered during cue presentation at 3, 6.5, and 9s), and six non-reinforced trials of cue D (no pellets delivered), presented in pseudo-random trial order.
3. Probe test: After conditioning, rats received 2 days of non-reinforced probe tests i.e. in extinction. On the first day, the probe test included a total of 6 trials of cue A and 6 trials of cue C, presented in alternating blocks of three trials of each stimulus (order counterbalanced across rats). On the second day, the probe test included a total of 6 trials of cue B (non-reinforced) and 6 trials of cue C, presented in pseudorandom order. The order of the probe test sessions was fixed for all rats (testing A/C on day 1, and B/D on day 2).

### Exclusion Criteria

Rats were excluded if they developed health issues during the experiment (Female n = 4, Male n = 5), if their catheter lost patency during SA training for cocaine and/or if they received fewer than 140 total infusions (cocaine or sucrose) over the 14 days of SA training (Female n = 7, male n = 4). Final numbers included in each cohort were: Cohort 1 (n = 5 Female, n = 6 Male), Cohort 2 (n = 4 Female, n = 6 Male), Cohort 3 (n = 6 Female, 5 Male), Cohort 4 (n = 12 Female, n = 12 Male), Cohort 5 (n = 13 Female, 15 Male). Final group numbers were: Suc_Veh N = 18 (n = 7 Female, n = 11 Male), Suc_D3a N = 15 (n = 6 Female, n = 9 Male), Coc_Veh N = 14 (n = 6 Female, n = 8 Male), Coc_D3a N = 17 (n = 10 Female, n = 7 Male).

The study aimed to include approximately n = 16 rats per group to achieve significant statistical power to detect the predicted treatment effect (based on pilot data, and published effect sizes using similar parameters (Jones et al., 2012; Wied et al., 2013)). This was achieved after training 5 cohorts. Analysis of the excluded animals did not reveal any systematic bias towards group membership or sex.

### Data analysis

#### Self-Administration

During SA training, the primary measures were the total number of lever presses on the active lever, inactive lever, and the total number of infusions per session. To account for differences in the number of reinforcers available, data are expressed as a percentage of the maximum available number of reinforcers i.e. for each rat/session Percent Max Response = (Total Responses/60)*100 for all animals, except Sucrose SA rats in cohort 1 where Percent Max Response = (Total Responses/300)*100.

#### Sensory Preconditioning

The primary response measure for SPC was the mean time spent in the food magazine during the auditory stimuli (CS), relative to baseline (PreCS, i.e. CS -PreCS). The baseline period was defined as the 10s immediately before stimulus presentation on each trial and was calculated separately for each corresponding stimulus/trial type. During the probe test, only the first 4 presentations of each stimulus were analyzed as this test was in extinction and responding rapidly declined during these sessions (however, this did not change the reported pattern of results).

Analysis of each stage was conducted using a linear mixed effects model ANOVA that included the following factors where relevant: between-subjects factors of SA (Suc, Coc), Treatment (Veh, D3a), Sex (M, F), and within-subjects factors of Cue (AB, CD), Stimulus (S1, S2), and Session (1-6). Significant interaction effects were explained by follow up analyses of any significant differences in lower order interactions, effects, or simple effects. Rates of acquisition across sessions were analyzed using linear trend contrasts from a set of planned orthogonal polynomial contrasts. When relevant, a Holm-Šídák correction was used to determine the significance of simple effects to control the nominal family-wise error rate at α = .05.

All analyses were reported from full models that included Sex as a factor. Sex differences are reported fully in supplementary material; however, any observed significant sex differences were transient and were not observed by the end of conditioning in stage 2, or during the critical probe tests. Cohort number was included as a random effect in all analyses but was removed as it did not account for any variance in the data. For all linear mixed effects models, a maximal random effects structure was used for all repeated/within-subjects measures and systematically simplified to deal with non-convergence (Barr et al., 2013; Meteyard & Davies, 2020), and Satterthwaite approximation was used to calculate degrees of freedom.

For the probe test, two additional analyses were used to test how the SA and Treatment groups were solving the SPC task, and whether these solutions were similar between groups. These were (1) Correlations between S1 and S2 stimulus pairs, and (2) a behavioral similarity analysis.

#### Correlations between S1 and S2 stimulus pairs

This analysis tested whether responding to cue A was predicted by B, and whether responding to cue C was predicted by D, and whether the strength of these relationships differed between each of the SA and Treatment groups. To achieve these simultaneous comparisons, a linear model was used to predict responding to S1 stimuli (A and C) with responding to S2 stimuli (B and D, respectively). The model included S1 as the outcome variable predicted by S2 x SA x Treatment x Cue x Sex (full factorial model, type III sums of square). S1 and S2 variables were continuous magazine duration (CS – PreCS) scores that were mean-centered within each cue. Sex did not contribute to the final model (main effects or interactions) and was dropped as a predictor from the final model. Follow up analyses to explain the higher order interaction were run separately for cue pairs AB and CD.

#### Behavioural similarity analyses

To quantify the full pattern of behavior both between and within cues A-D, a behavioral similarity analysis was used (similar to representational similarity analyses (Kriegeskorte et al., 2008). To account for within cue behavior, responding was first separated into 5s time bins within each 10s cue. For each group, a Pearson cross-correlation matrix of all cues/time bins was calculated to generate a behavioral similarity matrix. These matrices were then compared to test the similarity of the behavioral solutions between groups. This behavioral similarity analysis was conducted with Spearman rank correlations on the lower triangle of each group cross-correlation matrix. a Holm-Šídák multiple comparison correction was used to control the family-wise error rate at α = .05.

#### Statistical Software

All analyses were conducted in RStudio (Posit team, 2023; R Core Team, 2023). For specific analyses, the following packages were used: linear models -lm(), correlations -cor(), cor.test(), family-wise error rate correction -p.adjust (stats); Type III sums of squares for linear models, Anova() (car -Fox & Weisberg, 2019); Linear mixed effects models -mixed() (afex -Singmann et al., 2023); Simple effects – emmeans() (emmeans -Lenth, 2023).

## Results

### Self-Administration

Rats successfully acquired SA behavior in both the sucrose and cocaine groups (Figure 1B, C). Overall, responding on the active lever significantly increased over sessions, but not on the inactive lever (significant main effect of Session, *F*(13,106.46)= 9.18, *p* < .001; Lever, *F* (1,60.07)=98.61, *p* < .001; and Lever x Session interaction, *F* (13,112.30) =13.11, *p* < .001; positive linear trend for the Active Lever,*t* (264.63)=9.77,*p* < 001; linear trend for the Inactive Lever,*t* (264.63)=0.18, *p*=.854). Acquisition of sucrose SA responding was faster than cocaine SA (main effect of SA, *F* (1,60.08)=54.39,*p*.< .001; SA x Lever interaction,*F* (1,60.07)= 20.98, *p* < .001; SA x Lever x Session interaction,*F* (13,112.30)= 3.17, *p* < .001). Active lever responding increased significantly more quickly for the sucrose than the cocaine group (linear trend, Sucrose: Active Lever, *t* (263.89)= 9.20, *p* < .001; Cocaine: Active Lever,*t* (265.33)= 4.66, *p* < .001; significant SA x Session linear trend interaction on Active Lever, *t* (264.63)= 9.77, *p* < .001); inactive lever responding did not differ (linear trend, Sucrose: Inactive Lever, *t* (263.89)= 0.66, *p* .513; Cocaine: Inactive Lever, *t* (265.33)= − 0.39, *p* .= 699; non-significant SA x Session linear trend interaction on Inactive Lever, *t* (264.63)= 0.18, *p* .= 854). Importantly, SA acquisition was similar for animals assigned to the Vehicle and D3a treatment in the next stage of the experiment (analysis including Treatment as a factor; main effects and interactions with Treatment, all p > .118).

**Figure 1.**
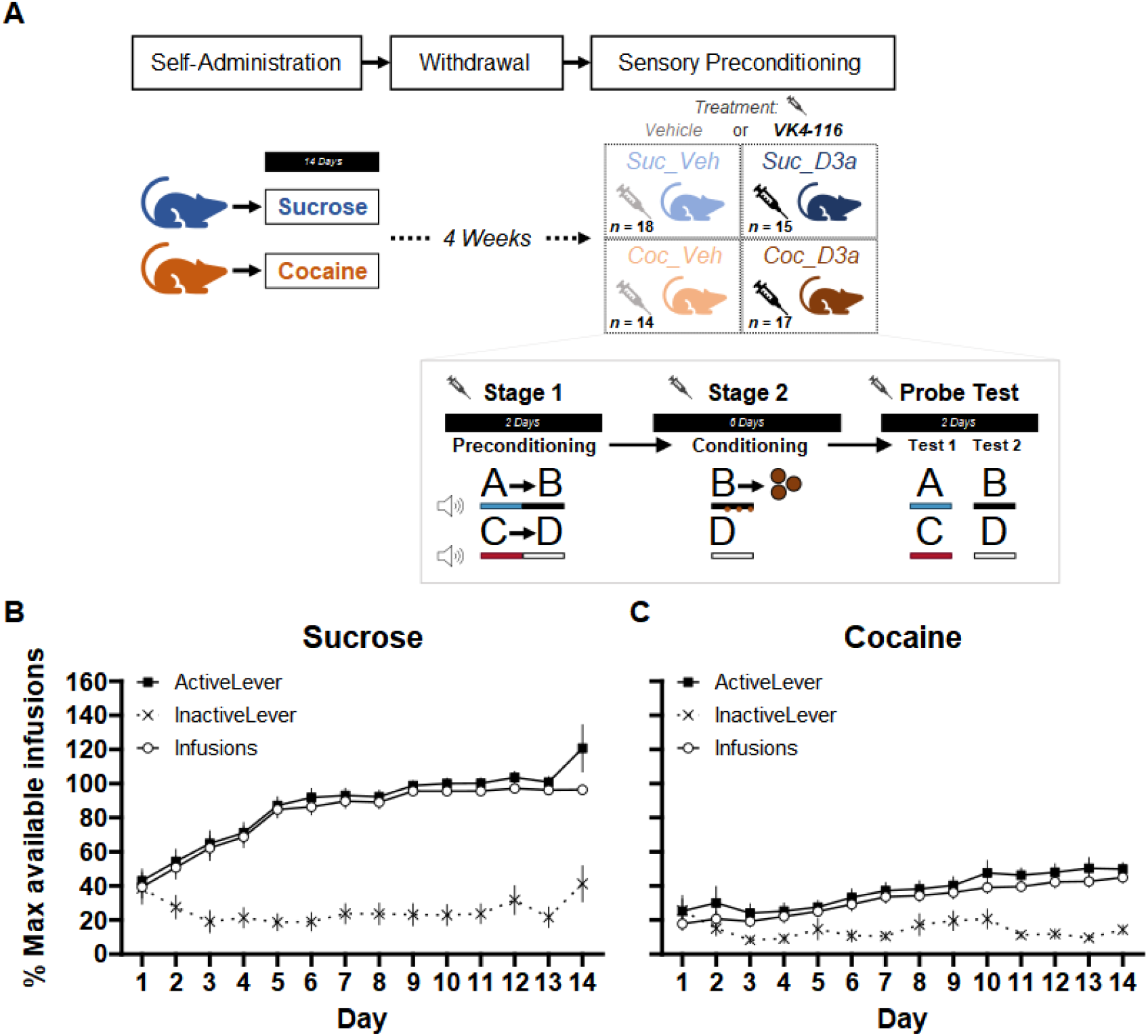
**(A)** *Timeline of experimental procedures:* (top) Rats first performed self-administration training (SA), followed by 4 weeks of withdrawal at home cages, and finally tested on a sensory-preconditioning (SPC) paradigm. Two separate groups of rats performed either sucrose, or cocaine, SA for 14 days. Following withdrawal, approximately half the animals in each SA group were assigned to a Treatment group and injected (i.p.) with either Vehicle or the D3-R antagonist VK4-116 (15 mg/kg) 30 mins before each session of the SPC paradigm. This created four separate SA x Treatment groups: Suc_Veh, Coc_Veh Suc_D3a, Coc_D3a (final group numbers indicated). *Design of the sensory preconditioning paradigm:* (inset) In preconditioning (stage **1**), rats were exposed to the sequential relationship between pairs of auditory stimuli A->B and C->D on separate trials. This was followed by conditioning (stage 2), where cue B was reinforced (3x sucrose pellets delivered during cue B presentation) whereas cue D was not reinforced. Finally, rats received two probe tests where cues A and C (test 1) or cues B and D (test 2; fixed test order) in extinction i.e. non-reinforced. The SPC effect: Greater anticipatory responding to A than C during the probe test indicates integration of learning across stages 1 and 2 i.e. A->B->Pellet, C->D->No-Outcome. The conditioning effect: greater anticipatory responding to cue B than D during the probe test indicates successful discriminative Pavlovian conditioning during stage 2. This extinction probe removes the pellet consumption responses that co-occur with anticipatory responding to the reinforced cue B during stage 2 conditioning. *Self-Administration training:* Rats were first trained on self-administration (SA) to press the active lever for either 10% sucrose liquid or cocaine (0.75/mg/kg/infusion). Rats in both the **(B)** sucrose and **(C)** cocaine groups successfully increased responding on the active lever, but not the inactive lever over 14 days of training. Overall, SA was acquired faster in the Sucrose group. Lever responses and reinforcer infusions are presented as a percentage of the maximum number of available infusions per session (max infusions was 60 for most animals; see methods for details). Error bars depict mean +/-SEM.

### Stage 1 -Preconditioning

Overall, responding during exposure to the non-reinforced cue pairs in preconditioning was low (Figure 2). Sucrose animals spent more time in the port than Cocaine SA animals (main effect of SA: Suc > Coc, *F* (1,56) = 6.69, *p* .=012; no main effect of Treatment: Veh vs D3a, *F* (1,56) 2.60, *p* .=113; no SA x Treatment interaction, *F* (1,56) 0.65, *p* =.422). Responding was also higher to the first stimulus in each pair (main effect of Stimulus: S1 > S2, *F* (1,56) = 11.09, *p* .=002), and slightly higher to the AB than the CD cues (main effect of Cue: AB > CD, *F* (1,56) = 4.19, *p* .=045; but no Cue x Stimulus interaction, *F* (1,56) = 0.06, *p* =.811). Importantly, there were no significant interactions between SA, Treatment, and Stimulus order or Cue pairs (all p >.022).

**Figure 2.**
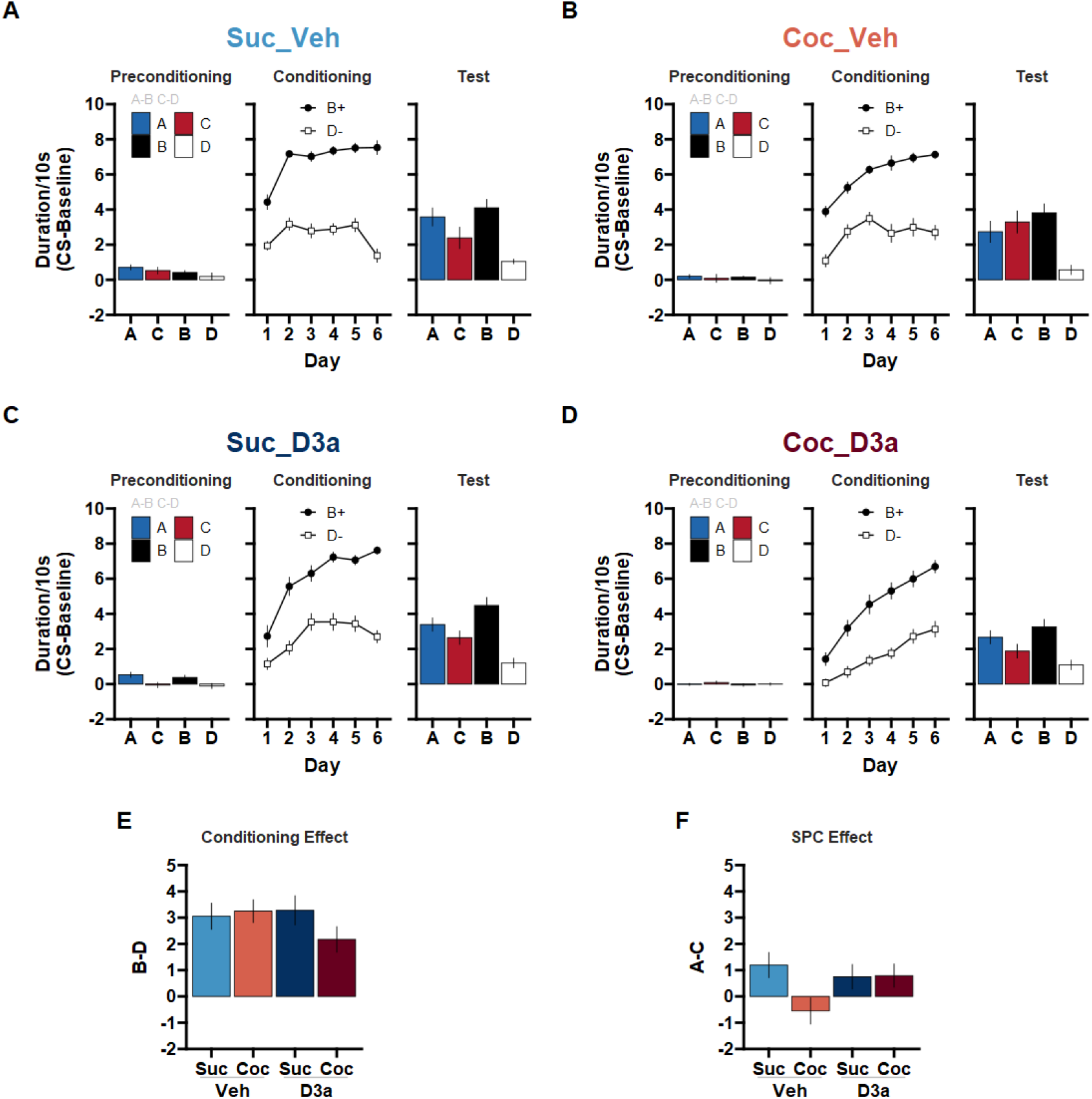
A history of cocaine self-administration disrupts sensory preconditioning (SPC) in rats treated with vehicle prior training sessions. The D3-R antagonist VK4-116 effectively treats sensory-preconditioning deficits in rats with a history of cocaine self-administration. Following SA training and 4 weeks of withdrawal, rats were trained on the SPC task. Prior to each session, rats were treated with either Vehicle or the D3-R antagonist VK4-116 (15 mg/kg; i.p.). **(A)** Suc_Veh: Sucrose SA rats given Vehicle. **(B)** Coc_Veh: Cocaine SA rats given Vehicle. **(C)** Suc_D3a: Sucrose SA rats given VK4-116. **(D)** Coc_D3a: Cocaine SA rats given VK4-116. Stage 1 -Preconditioning (A->B, C->D), Stage 2 -Conditioning (B->Pellets, D->No outcome), and Probe test (A, C, B, D: in extinction) behavior are presented for each group (left, middle, right). Discriminative responding to the cues is depicted as the duration of time spent in the food cup during the CS, above the pre-CS baseline (mean +/-SEM). **(E)** The Conditioning effect: The difference in responding to cues B and D during the probe test provides an index of the conditioning effect such that scores above 0 reflect successful conditioning. **(F)** The SPC effect: The difference in responding to cues A and C during the probe test provides an index of the SPC effect such that scores above 0 reflect successful SPC. Difference scores were calculated as the difference in discriminative responding to each cue from the Probe test in A-D (mean +/-SEM).

### Stage 2 -Conditioning

All four groups successfully increased responding to cue B more than cue D by the end of conditioning (main effect of Cue, *F* (1,56) = 726.65, *p* < .001; Session, *F* (5,560) = 79.20, *p* < .001; and significant Cue x Session, *F* (5,560) =13.42, *p* < .001). However, it is important to note that pellets were delivered during presentations of cue B, and responding in this stage reflects both anticipation and consumption of the pellets. Responding to cues B and D during the probe test (i.e. in extinction) can provide a test of any differences in anticipatory responding without this confound.

There was greater cue discrimination (B > D) in the sucrose than the cocaine SA groups (SA x Cue interaction, *F* (1,56) = 6.35, *p* .015; significant cue discrimination in both SA conditions, Sucrose: B > D, *t* (56) = 21.08, *p* < .001; Cocaine: B > D, *t* (56) = 17.09, *p* < .001), however there was also a greater linear increase in responding to both cues in the cocaine than sucrose groups (SA x Session interaction, *F* (5,560) = 5.39, *p* < .001; significant positive linear trend in both SA conditions: Sucrose, *t* (560) = 9.60, *p* < .001; Cocaine, *t* (560) = 14.92, *p* < .001; Overall responding main effect of SA: Suc > Coc, *F* (1,56) =14.42, *p* < .001).

When looking at the effects of Treatment groups, there was greater cue discrimination (B > D) in the Vehicle than the D3a groups (Treatment x Cue interaction, *F* (1,56) = 8.21, *p* .006; significant cue discrimination in both Treatment conditions, Vehicle: B > D, *t* (56) = 21.04, *p* < .001; D3a: B > D, *t* (56) = 17.07, *p* < .001), however there was also greater linear increase in responding to all cues in the D3a than vehicle Treatment groups (Treatment x Session interaction, *F* (5,560) = 11.27, *p* < .001; significant positive linear trend in both SA conditions, Vehicle: linear trend, *t* (560) = 7.19, *p* < .001; D3a: linear trend, *t* (560) = 17.41, *p* < .001; and an overall main effect of Treatment: Veh > D3a, *F* (1,56) =12.40, *p* .001).

Overall, the effects of SA and Treatment did not interact significantly (SA x Treatment and higher order interactions with Cue and Session, p > .054).

#### Final Session

Focusing on the final day of conditioning (Session 6), all groups responded significantly more to cue B than D (main effect of Cue, *F* (1,56) =420.36, *p* < .001). Between groups, there were no differences in responding to cue B, however responding to cue D was higher in the Cocaine than Sucrose SA groups (SA x Cue interaction, *F* (1,56) = 11.15, *p* .002; Cue B: Suc vs Coc, *t* (102.22) = −1.47, *p* .144; Cue D: Suc < Coc, *t* (102.22) = 2.45, *p* .016), and greater in the D3a than the Vehicle Treatment groups (Treatment x Cue interaction, *F* (1,56) = 6.45, *p* .014; Cue B: Veh vs D3a, *t* (102.22) = −0.37, *p* .714; Cue D: Veh < D3a, *t* (102.22) = 2.62, *p* .010). All remaining main effects and interactions between SA, Treatment, Cue, and Sex failed to reach significance (p > .100).

### Stage 3 -Probe Test

During the probe test (Figure 2A-D), all groups demonstrated successful stage 2 conditioning (i.e., the conditioning effect, B > D; Figure 2E). In contrast, all groups except for the untreated cocaine group (Coc_Veh), responded more to cue A than C (i.e. the SPC effect, A > C; Figure 2F). This pattern of group differences was supported by a significant SA x Treatment x Cue x Stimulus interaction (*F* (1,112) = 5.90, *p* .017; a main effect of Cue pair, AB > CD, *F* (1,112) = 89.16, *p* < .001; and a Cue x Stimulus interaction, *F* (1,112) = 41.78, *p* < .001; all remaining effects, including sex differences did not reach significance, p > .098), and explored with separate follow-up analyses within each Treatment condition.

#### Vehicle Treatment

In the Vehicle treated groups, sucrose SA rats showed a significant SPC effect that was abolished in cocaine SA rats. This was supported by a significant SA x Cue x Stimulus interaction (*F* (1,56) =5.01, *p*= .029), such that the SPC effect (A > C) was significant in Suc_Veh (Suc_Veh: A > C, *t* (56) = 2.58, *p*= .013) but not Coc_Veh rats (Coc_Veh: A vs C, *t* (56) = 1.05, *p*= .297; reflecting a significant SA x Cue for S1 stimuli, S1: SA x Cue *t* (56) = −2.51, *p*= .015). Importantly, both groups showed significant evidence of successful discriminative responding to the previously rewarded cue B compared to the non-reinforced cue D (Suc_Veh: B > D, *t* (56) = 5.88, *p* < .001; Coc_Veh: B > D, *t* (56) = −1.05, *p*= .297), and this effect was of similar magnitude in both groups (no SA x Cue interaction for S2 stimuli, S2: SA x Cue t 56 0.66, *p*= .512). This finding successfully replicates our earlier report that a history of cocaine SA disrupts SPC in rats, and it further extends this by suggesting that there are no significant sex differences in this effect.

#### D3a Treatment

In D3a treated groups, the SPC effect was significant and did not differ between sucrose and cocaine SA rats. This was supported by a significant Cue x Stimulus interaction (*F* (1,28) = 15.25, *p* =.001; and significant main effect of Cue pair, AB > CD, *F* (1,28) = 42.89, *p* < .001) that did not differ between Suc_D3a and Coc_D3a groups (no Treatment x Cue x Stimulus interaction, *F* (1,28) = 1.55, *p*= .224). This reflected a significant SPC effect across both Suc_D3a and Coc_D3a groups (A > C, *t* (55.90) = 2.02, *p* = .048), as well as a significant, albeit larger, effect of discriminative responding (B > D, *t* (55.90) = 7.43, *p* < .001). Notably, both of these effects were of similar magnitude in both groups (no SA x Cue interaction for S1 or S2 stimuli; SPC effect, S1: SA x Cue, *t* (55.90) = 0.26, *p* = .797; conditioning effect, S2: SA x Cue, *t* (55.90) = −1.46, *p* = .149). Therefore, treating cocaine-experienced rats with the D3-receptor antagonist VK4-116 successfully recovered deficits in SPC.

### Group differences in SPC task interpretation

Although no cues were used in our self-administration sessions, it is nevertheless plausible that different histories of cocaine or sucrose SA (i.e. Suc_Veh and Coc_Veh) could change the nature of how the cues are represented, integrated, or generalized across the stages of the SPC task, i.e. how the task is “solved”. In control animals, the SPC effect can be supported by a number of mechanisms that are likely to be a parameter dependent (Holmes et al., 2022; Killcross & Blundell, 2002). In the present version of the task, the predominant mechanism in controls is likely to be associative inference during the probe test (Hart et al., 2020; Sharpe et al., 2017) i.e. at test, responding to cue A is driven by recall of the A->B (stage 1) and B->Pellet (stage 2) associations that are integrated to infer that A->B->Pellet. This is suggested by several lines of evidence, including the OFC-dependence of responding in the probe test (Jones et al., 2012; Sadacca et al., 2018), as well as the failure of the pre-conditioned cue to support conditioned reinforcement (Sharpe et al., 2017), both of which are contrary to other mechanisms, especially mediated learning, in which value would accrue directly to the preconditioned cue (Killcross & Blundell, 2002). This solution to SPC is disrupted by a history of cocaine use, both here in the Coc_Veh group as well as in our prior study (Wied et al., 2013). If VK4-116 (D3a) is successfully “treating” the effect of Cocaine SA, then the Coc_D3a group should also have a task solution that is more similar to the Suc_Veh than the Coc_Veh groups. We tested the similarity of the group solutions using two complementary approaches: (1) testing specific predictions about the A∼B and C∼D relationships in each group, and (2) comparing the similarity of the overall task solution between groups.

To test this, we first examined probe test behavior to see if responding to cues A and C was related to the level of conditioning to cues B and D, i.e. is there an S1->S2 relationship? Specifically, in the Suc_Veh group we expected a strong positive correlation between learning about reinforced cue B and responding to cue A, and no correlation between non-reinforced cue D and cue C during the probe test. In contrast, in the Coc_Veh group we expected that responding would not be correlated between cues pairs A-B and C-D. Importantly, we predicted that the D3a treatment in the Coc_D3a group would recover the strong positive correlation between responding to cues A and B, but not C and D. The pattern of group differences in slopes between cues A∼B and C∼D was compared by fitting a linear model predicting responding to the first stimulus in each pair (S1: A/C) with responding to the relevant second cue in each pair (S2: B/D), as well as categorical factors of Cue pair (AB/CD), Treatment, and SA.

Consistent with our predictions (Figure 3A), there was a significant positive correlation between A and B in the Suc_Veh and Coc_D3a groups but not in the Coc_Veh group or, unexpectedly, in the Suc_D3a group (Suc_Veh: A ∼ B, b = 0.75, 95% CI [0.35,1.15], *t* (16) = 3.96, *p* = .001; Coc_D3a: A ∼ B, b = 0.57, 95% CI [0.20,0.95], *t* (15) = 3.26, *p* = .005; Coc_Veh: A ∼ B,= b 0.09, 95% CI [−0.66,0.83], *t* (12) =0.25, *p* = .805; Suc_D3a: A ∼ B, b = −0.22, 95% CI [−0.70,0.26], *t* (13) = 0.99, *p* = 340). These slope differences between SA and Treatment groups was supported by a significant SA x Treatment interaction (A ∼ B: SA x Treatment, *F* (1,56) = 9.68, p 003), which reflected significant differences between slopes for Coc_Veh and Suc_Veh groups (A ∼ B: Coc_Veh < Suc_Veh, *t* (56) = −2.10, *p* = .040), and Coc_D3a and Suc_D3a groups (A ∼ B: Coc_D3a > Suc_D3a, *t* (56) = 2.30, *p* = .025).

**Figure 3.**
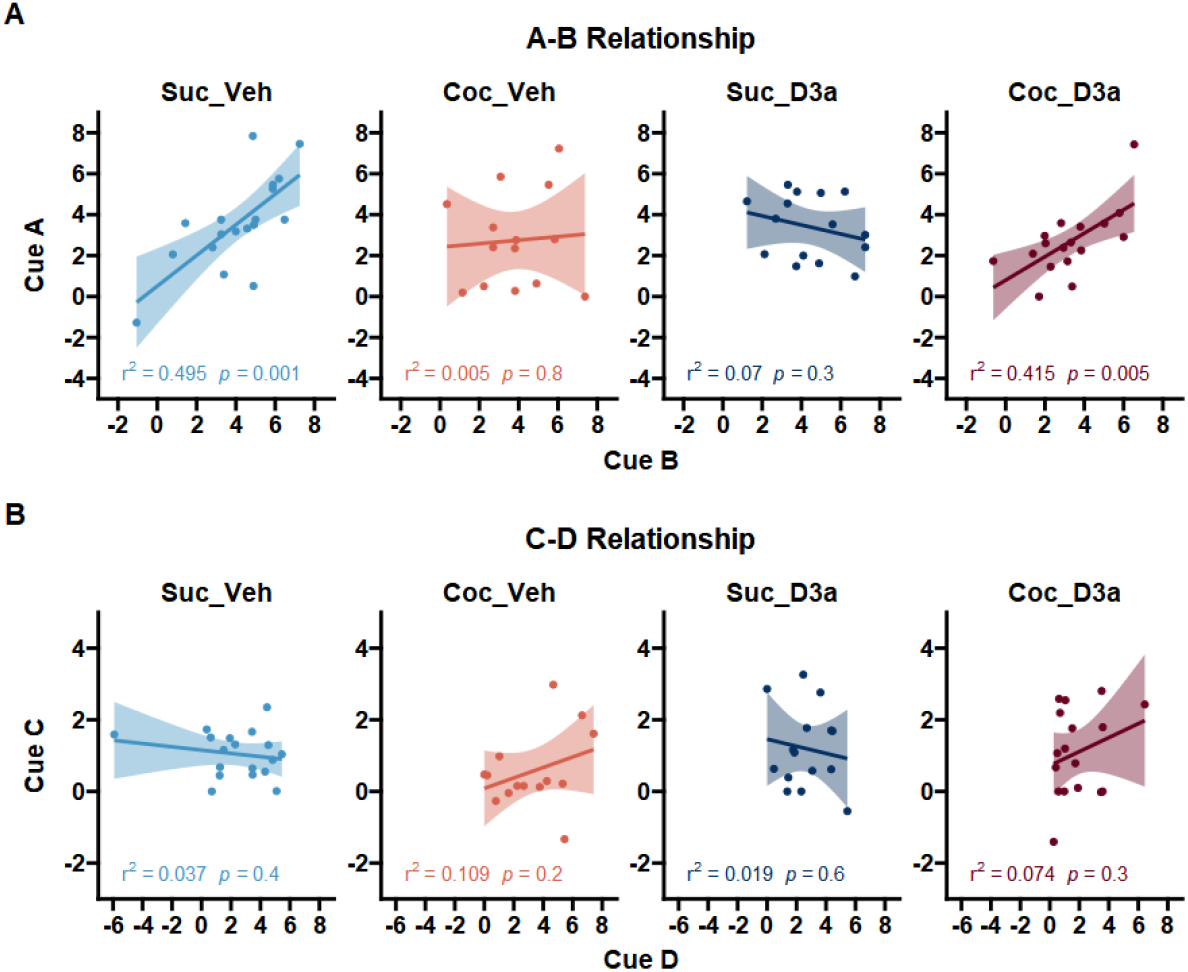
Testing whether probe test responding reflects the relationships between the cue pairs that were presented during stage 1 preconditioning and stage 2 conditioning (A->B->Pellet, C->D->No-outcome). A *A-B relationship*: Responding to cue A is driven by learning to B in both the control group (Suc_Veh) and treated cocaine group (Coc_D3a), but not in the untreated cocaine (Coc_Veh) and treated control groups (Suc_D3a). B *C-D relationship*: There was no relationship between responding to cue C and D. Data points indicate probe test responding for individual subjects for the first cue (y-axis) and second cue (x-axis) in each pair, and plotted separately for each group (panels from left to right). Responding was defined as the duration of time spent in the food cup during the CS, above the pre-CS baseline. Lines and error shading from linear model fit, and corresponding model r^2 and p-values presented at the bottom of each plot.

**Figure 4.**
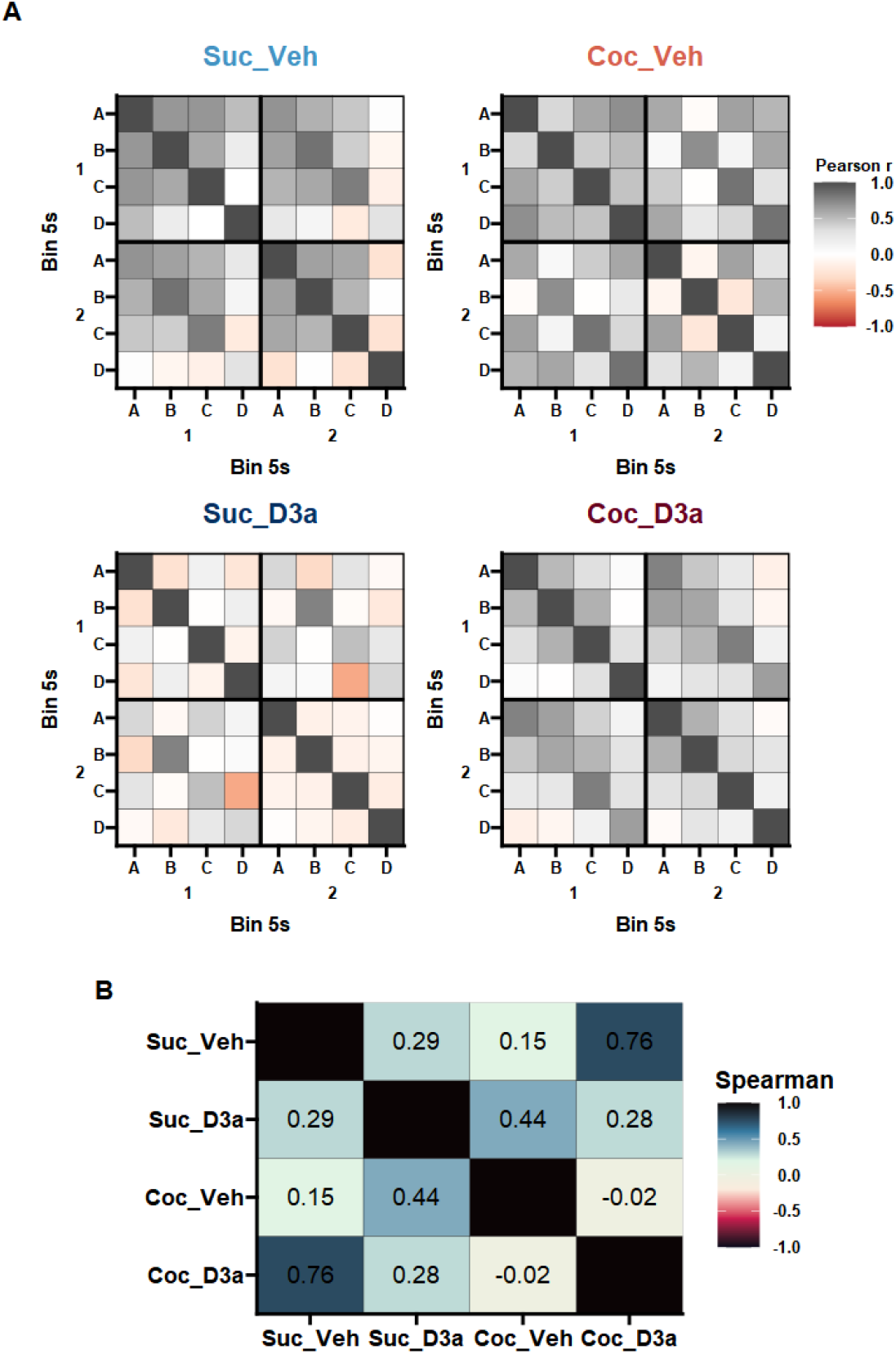
Behavioral similarity analysis of probe test responding. **A** Behavioral similarity matrices for Sucrose (left) or Cocaine (right) SA groups, and Vehicle (top) or D_3_R antagonist (bottom) treatment groups. For each group, the similarity of responding between and within cues was quantified by separating responses within each cue into 5s time bins (Supplementary Figure 1) and generating a cross correlation matrix between all the resultant cue epochs. Color values are plotted to represent the correlation values (Pearson’s r), to provide a visual summary of the group specific pattern of response relationships. The behavioral similarity patterns were similar between the untreated control group (Suc_Veh) and the cocaine group treated with the D_3_R antagonist (Coc_D3a), but different to the untreated cocaine group (Coc_Veh) and the control group treated with the D_3_R antagonist (Suc_D3a). **B** *Behavioral similarity analysis*: A Spearman correlation was used to test the similarity between the group similarity matrices. Numbers (and corresponding color values) indicate the rank correlation (Spearman’s rho) between the lower diagonal of the cross-correlation matrices above. An analysis of just the between-cue similarities (i.e. not considering within-cue time bins) supported similar conclusions (Supplementary Figure 2).

In contrast to the A∼B correlations, there were no significant relationships between cue C and D in any group (Figure 3B; Suc_Veh: C ∼ D,b = − 0.81, 95% CI [3.01,1.38], *t* (16) = − 0.79, *p* = .443; Coc_Veh: C ∼ D,b = 0.74, 95% CI [− 0.59,2.08], *t* (12) = 1.21, *p* =.249; Suc_D3a: C ∼ D,b = 0.19, 95% CI [1.01,0.63], *t* (13) = − 0.50, *p* =.625; Coc_D3a: C ∼ D, b = 0.38, 95% CI [− 0.36,1.11], *t* (15) = 1.09, *p* = 292). The lack of significant group differences in the C-D relationship was consistent with a non-significant SA x Treatment interaction (C ∼ D: SA x Treatment, 1,56 0.68, *p*= .414; all remaining main effects and interactions with cue D, p > 399). Finally, this overall pattern of group differences in slopes between cues A∼B but not C∼D was supported by a significant SA x Treatment x Cue x S2 interaction (*F*(1,112)= .4.06, *p*= .046). These results supported our predictions that responding to cue A was uniquely related to learning about cue B in control rats (Suc_Veh), which was disrupted by a history of cocaine use (Coc_Veh) but successfully recovered by D3a treatment (Coc_D3a). Surprisingly, the D3a treatment disrupted the A∼B relationship in sucrose control rats (Suc_D3a) but not the SPC effect, suggesting that these rats were employing a different strategy to solve the SPC task.

### Behavioral Similarity

Next, we tested the similarity of the overall task solutions in each group using a behavioral similarity analysis approach. How a task is solved is likely to be reflected in the pattern of multivariate relationships both between and within each cue. To capture the pattern of responding within each cue, the 10s cue period was split into early and late epochs (i.e. 5s bins), a standard approach (Delamater & Holland, 2008; Gallagher et al., 1999; Holland, 1977) that also reflected the observed pattern of responding within cue (Supplementary Figure 1). A Pearson cross-correlation matrix was calculated across all 8 (cue[4] x time[2]) epochs to create a behavioral similarity matrix for each group (Figure 3). Evidence of significant similarity was found between Suc_Veh and Coc_D3a groups (Spearman’s rank correlation, Suc_Veh vs Coc_D3a, *rs*= .76, *S* = 894.00, *p* < .001), but not between the other groups (Suc_Veh vs Coc_Veh, .*r*_*s*_= .15, *S* =3,088.00, *p* .> 999; Suc_Veh vs Suc_D3a, *r*_*s*_= .29, *S* = 2,594.00, *p* = .805; Coc_Veh vs Coc_D3a, .*r*_*s*_= .−02, *S* = 3,716.00, *p* >.999; Coc_Veh vs Suc_D3a, *r*_*s*_= .44, *S* = 2,052.00, *p* = .123; Coc_D3a vs Suc_D3a, *r*_*s*_= .28, *S* =2,622.00, *p* = .871; Holm-Sidak corrected). These findings suggest that the D3a treatment in cocaine-experienced rats was able to use a similar solution to the SPC task as the vehicle treated sucrose control rats. Furthermore, in sucrose experienced rats the D3a treatment did not disrupt the SPC effect but has changed how they are solving the task relative to the other groups.

## Discussion

A history of cocaine use in humans and other animals is associated with behavioral inflexibility linked an increase in mesolimbic D_3_R availability, and a lack of behavioral insight, i.e., an inability to use learned information to mentally simulate and make inferences about cause and effect. The present study tested whether the novel selective D_3_R-antagonist, VK4-116, could effectively treat impaired behavioral inferences in sensory preconditioning (SPC) in rats following withdrawal from cocaine self-administration.

We found that, a history of cocaine-self administration caused a significant impairment in the ability of vehicle-treated rats to make behavioral inferences by integrating prior knowledge about the relationships between cues and outcomes in the SPC task. However, treating cocaine-experienced rats with the D_3_R-antagonist VK4-116 effectively restored behavioral inference in a manner strikingly similar to vehicle-treated sucrose-control rats. This was evident in two metrics of responding to cue A and C at test that reflect the inference that A->B->Reward and C->D->Nothing: (1) responding more to cue A than C, and (2) the strength of responding to cue A reflected the strength of responding to cue B. Surprisingly, VK4-116 also partially disrupted behavioral inference in sucrose-experienced control rats such that while responding to A was greater than C, it was unrelated to responding to cue B. Importantly, these cocaine and VK4-116 treatment effects were not simply due to differences in learning or responding to cues B and D, i.e. non-inference based behavior.

Overall, these findings suggest that both VK4-116 and cocaine-history shifted how the SPC task was interpreted. This was tested with a behavioral similarity analysis, which confirmed that VK4-116 treated cocaine rats were solving the task in a manner similar to vehicle-treated sucrose control rats, but not similar to untreated cocaine rats or VK4-116 treated sucrose rats. These data provide evidence that D_3_R play a key role in the loss of behavioral insight and inference processes following cocaine use and suggest VK4-116 as a promising pharmacotherapy to selectively treat these symptoms in CUD.

### D_3_R function is related to impaired behavioral inference following cocaine use

The selective D_3_R-antagonist VK4-116 impaired behavioral inference in drug-naïve rats but effectively restored it in rats after prior-cocaine exposure that has previously been shown to increase brain D_3_R availability (Groman et al., 2020). This bidirectional effect of VK4-116 suggests that D_3_R plays a causal role in modulating behavioral inference, and that increased midbrain D_3_R availability is a key factor in impaired behavioral inference following cocaine use. This account is also consistent with the extant literature on the role of D_3_R in cocaine use and its behavioral sequalae (Newman et al., 2023). In drug-naïve rodents, higher levels of midbrain D_3_R availability predicts behavioral inflexibility in probabilistic reversal learning tasks, as well as increased motivation and escalation of subsequent cocaine SA (Groman et al., 2016, 2020). Furthermore, pharmacological activation of D_3_R with the D_3_R-preferring agonist pramipexole, in drug-naïve rats, exacerbates probabilistic reversal learning deficits in a manner similar to cocaine use (Groman et al., 2016). While there is evidence that individuals with high levels of D_3_R availability are more likely to use cocaine, subjects with cocaine experience exhibited increased D_3_R availability in the striatum and midbrain in humans and non-human primates (Boileau et al., 2012; Matuskey et al., 2014; Payer et al., 2014; Segal et al., 1997; Staley et al., 1994; Staley & Mash, 1996). In the present study, we found that pretreatment with a selective D3R antagonist (VK4-116) restored cocaine use-induced deficit in behavioral inference and flexibility in rodents, suggesting that higher D3R availability in the brain could be a risk factor in the development of cocaine use disorder.

There also appears to be a non-monotonic relationship between D_3_R function and behavioral inference because blocking D_3_R function with VK4-116 in drug naïve rats also impaired behavioral inference. This would suggest that D_3_R may also functionally involve behavioral inference process in healthy subjects. Higher D_3_R availability in cocaine users is likely to account for the lack of individual differences in behavioral inference observed in both cocaine-treated rats with VK4-116 and drug-naïve rats.

### Relationship between D_3_R and orbitofrontal function in CUD

There appears to be a strong connection between midbrain D_3_R and OFC function. D_3_Rs are inhibitory G-protein coupled receptors. Activation of D3Rs in dopamine neurons in substantia nigra and ventral tegmental area inhibits dopamine release in striatum and OFC (Gurevich & Joyce, 1999; Heidbreder & Newman, 2010; Koob & Volkow, 2010; Newman et al., 2021; Sesack & Grace, 2010). Consistent with this relationship, imaging studies also indicate that OFC hypoactivity is a consistent feature in abstinent cocaine users, and is correlated with increased D_3_R availability in substantia nigra (Volkow et al., 1993; Volkow & Fowler, 2000; Wilcox et al., 2011). In non-clinical populations, midbrain D_3_R availability is also correlated with both resting state OFC activity, and the functional connectivity between OFC subregions and multiple key large-scale neural systems such as salience executive control networks, basal ganglia/limbic network, and the default mode network (Cole et al., 2012). Thus far, it is unknown which neuronal cell type displays D_3_R up-regulation in subjects with CUD. If it occurs mainly in midbrain DA neurons, blockade of abnormal D_3_Rs would disinhibit (increase) dopamine neuron activity, which may subsequently increase OFC neuronal activity and improve functional connectivity between the OFC and other networks. This may in part underlie the therapeutic effects of D_3_R antagonism on cocaine-induced impairment of behavioral inference.

Recent evidence from cocaine self-administration and forced abstinence studies in rodents and non-human primates also supports a causal link between cocaine use, increased brain D_3_R availability, OFC hypofunction, and behavioral deficits in tasks like probabilistic reversal learning (Groman et al., 2016, 2020; Jedema et al., 2021). Indeed, there is significant overlap between the behavioral deficits following cocaine use and OFC dysfunction. These deficits include such behavioral inflexibility in reversal learning, and impaired inferences across task states in SPC, Pavlovian over-expectation, and outcome devaluation (Schoenbaum & Shaham, 2008).

We have previously shown that cocaine experience disrupts OFC function and impairs insight-based behavior in rats, and optogenetic activation of pyramidal neurons in OFC can successfully restore insight-based behavior (Lucantonio et al., 2014). These findings parallel the present study and suggest that increased midbrain D_3_R availability and OFC hypofunction are correlated and underlie deficits in insight-based behavior following cocaine use. Treatment with the selective D_3_R-antagonist VK4-116 may well have its effect in the cocaine experienced rats here through restoring the normal balance in dopamine effects on OFC function altered by drug experience.

## Summary

Here we show the first evidence for the efficacy of VK4-116, a novel and highly selective D_3_R antagonist, in the treatment of impaired behavioral inferences, a persistent aspect of behavioral inflexibility caused by cocaine use. These findings support established evidence in rodents that D_3_R antagonists significantly reduce drug taking and relapse in rodent models of cocaine, nicotine, and oxycodone self-administration (Ross et al., 2007; Z.-X. Xi et al., 2005; You, Bi, Galaj, Kumar, Cao, Gadiano, Rais, Slusher, Gardner, Xi, & Newman, 2019). In contrast to antagonists that target D_2_Rs, VK4-116 does not appear to significantly disrupt overall activity levels or cause anhedonia (You, Bi, Galaj, Kumar, Cao, Gadiano, Rais, Slusher, Gardner, Xi, & Newman, 2019). Therefore, selective D_3_R antagonists such as VK4-116 are a promising therapeutic target for CUD and other SUDs (Groman et al., 2020; Newman et al., 2023).

## Acknowledgements

We thank J. Cao (Medicinal Chemistry Section, NIDA-IRP) for the synthesis of VK4-116. This work was supported by Z1A DA000424 (AHN and Z-X Xi) and Z1A DA000587 (GS) at the Intramural Research Program at the National Institute on Drug Abuse. The opinions expressed in this article are the authors’ own and do not reflect the view of the NIH/DHHS. The authors have no conflicts of interest to report.

## Data availability

Data available online at: https://osf.io/hpe4v/?view_only=bb9830485d394b188c79bb54aa826900

