## Supplemental Material for "The selective D3-Receptor antagonist VK4-116 effectively treats behavioral inflexibility in rats caused by self-administration and withdrawal from cocaine"

### Supplementary Analysis

#### Supplementary analysis 1: Self-Administration

***SA Sex differences:*** There were significant sex differences in sucrose but not cocaine SA (SA x Sex, $F\left( 1,60.08 \right)=4.21$, $p=.045$; SA x Lever x Sex, $F\left( 1,60.07 \right)=3.30$, $p=.074$; SA x Lever x Session x Sex, $F\left( 13,112.30 \right)=1.96$, $p=.031$). These sex differences were restricted to the active lever (SA x Sex x Session linear trend: Active Lever $t\left( 264.63 \right)=2.90$, $p=.004$; but not on the inactive lever, SA x Sex x Session linear trend: Inactive Lever, $t\left( 264.63 \right)=-0.61$, $p=.540$), and only for sucrose SA (Sex x Session linear trend: Active Lever - Sucrose, $t\left( 263.89 \right)=3.21$, $p=.001$; Sex x Session linear trend: Active Lever - Cocaine, $t\left( 265.33 \right)=-0.91$, $p=.365$). Specifically, while both sexes increased responding on the active lever, the rate of increase was significantly lower in females than males (Male: Sucrose, Active Lever - significant positive linear trend over Session, $t\left( 264.16 \right)=9.89$, $p<.001$; Female: Sucrose, Active Lever - significant positive linear trend over Session, $t\left( 263.71 \right)=3.85$, $p<.001$).

#### Supplementary analysis 2: Stage 1 - Preconditioning

***Sex differences:*** In general, there were no main effects or interactions with sex, except for a significant SA x Treatment x Cue x Sex interaction ($F\left( 1,56 \right)=5.51$, $p=.022$; all other p > .118). This reflected a sex difference in Suc_Veh group males that responded higher overall to the AB than the CD cues compared to Suc_Veh females, a pattern that was not significant in the other groups.

Specifically, this significant 4-way interaction reflected a significant 3-way SA x Cue x Sex interaction in the Veh ($F\left( 1,28 \right)=5.70$, $p=.024$) but not the D3a groups ($F\left( 1,28 \right)=0.45$, $p=.508$). In the Veh treatment groups, this significant 3-way interaction revealed a significant 2-way SA x Sex interaction in Sucrose but not Cocaine groups (Vehicle - Sucrose: SA x Sex, $t\left( 28 \right)=2.36$, $p=.026$; Vehicle - Cocaine: SA x Sex, $t\left( 28 \right)=-1.09$, $p=.284$), such that responding to Cue pair AB was greater than CD in Suc_Veh but not Coc_Veh females (Suc_Veh: Females - AB > CD, $t\left( 28 \right)=2.40$, $p=.023$, Coc_Veh: Females - AB vs CD, $t\left( 28 \right)=-0.42$, $p=.675$), but not males (Suc_Veh: Males - AB vs CD, $t\left( 28 \right)=-0.77$, $p=.447$, Coc_Veh: Males - AB vs CD, $t\left( 28 \right)=1.18$, $p=.248$).

#### Supplementary analysis 3: Stage 2 - Conditioning

***Sex Differences:*** There was a significant 5-way SA x Treatment x Sex x Cue x Session interaction ($F\left( 5,560 \right)=3.15$, $p=.008$), which was decomposed using interaction contrasts where Session was coded as polynomial linear trend contrast to quantify the rate of increase or decrease in responding over sessions. This 5-way interaction reflected significant sex differences in acquisition over sessions to cue B in Suc_Veh rats (linear trend contrast over sessions, Suc_Veh: Cue B, significant Sex x Session, $t\left( 560 \right)=2.91$, $p=.004$, Suc_Veh: Cue D, Sex x Session, $t\left( 560 \right)=-0.23$, $p=.817$; all remaining Sex x Session interactions for each cue and group, p > .067). Specifically, there was a significant linear increase in responding over sessions to cue B in the males but not the females (Males, $t\left( 560 \right)=6.72$, $p<.001$; Females, $t\left( 560 \right)=1.64$, $p=.102$). This was supported by a significant 3-way Sex x Cue x Session interaction only in the Suc_Veh group ($t\left( 560 \right)=-2.22$, $p=.027$; but not significant in the other groups, p > .256), and a significant 4-way SA x Sex x Cue x Session interaction for the Vehicle but not D3a groups (Vehicle, $t\left( 560 \right)=-1.86$, $p=.063$; D3a, $t\left( 560 \right)=0.23$, $p=.818$). However, since responding to cue B during the conditioning stage conflates anticipation and consumption of the pellets, it is hard to interpret the exact nature of these sex differences.

### Supplementary Figures

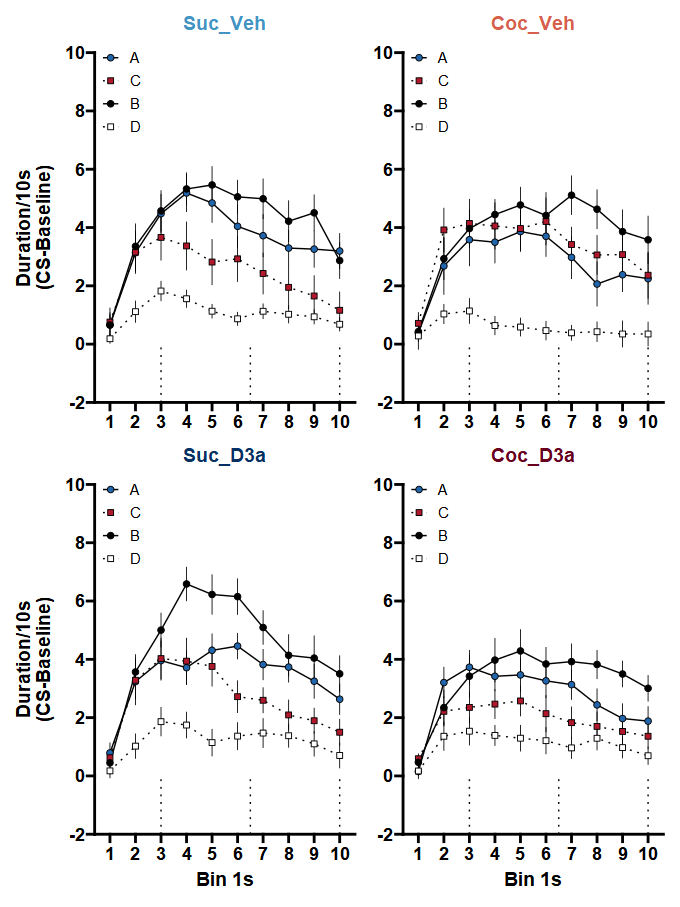
***Supplementary Figure 1.*** Within cue response patterns during the probe test for the Sucrose (left) or Cocaine (right) SA groups, and Vehicle (top) or D3-R antagonist (bottom) treatment groups. Responding in 1s time bins within each cue was calculated as the duration of time spent in the food cup during each time bin (as a rate per 10s), above the corresponding full 10s pre-CS baseline (mean +/- SEM). Dotted lines at 3, 6.5, and 10s indicate the time at which a pellet would have been delivered during reinforced presentations of cue B in stage 2 conditioning. The expected timing of pellet delivery is a within-cue response pattern that may reflect group specific solutions to the sensory preconditioning task.

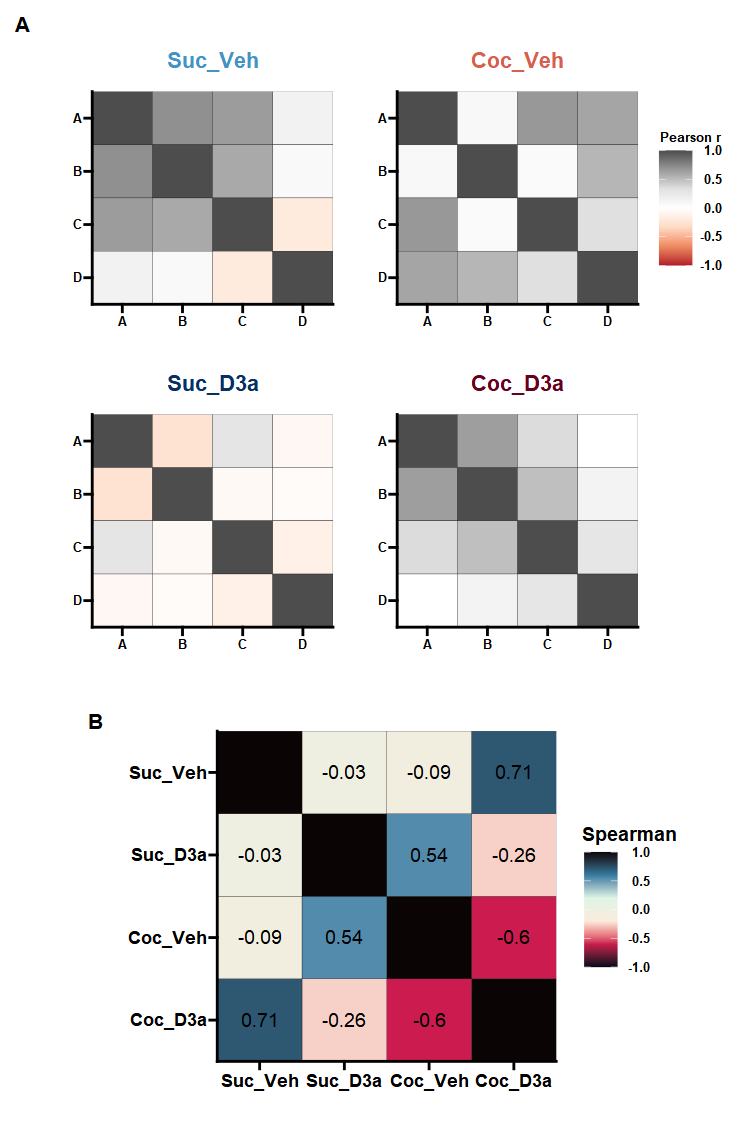
***Supplementary Figure 2.*** Behavioral similarity analysis of probe test responding between cues, but ignoring within-cue response patterns (see Figure 4). **A** Behavioral similarity matrices for Sucrose (left) or Cocaine (right) SA groups, and Vehicle (top) or D3-R antagonist (bottom) treatment groups. For each group, the similarity of responding between cues was quantified by generating a cross-correlation matrix between all the cues. Color values are plotted to represent the correlation values (Pearson’s r), to provide a visual summary of the group specific pattern of response relationships. The behavioral similarity patterns were similar between the untreated control group (Suc_Veh) and the cocaine group treated with the D3-R antagonist (Coc_D3a), but different to the untreated cocaine group (Coc_Veh) and the control group treated with the D3-R antagonist (Suc_D3a). **B** *Behavioral similarity analysis:* A Spearman correlation was used to test the similarity between the group similarity matrices. Numbers (and corresponding color values) indicate the rank correlation (Spearman’s rho) between the lower diagonal of the cross-correlation matrices above.
